# Cryo-EM structure of *Chlamydomonas* Photosystem I complexed with the alternative electron donor cytochrome *c*₆

**DOI:** 10.1101/2025.09.08.674899

**Authors:** Yu Ogawa, Gyana Prakash Mahapatra, Yuval Milrad, Michelle Schimpf, Genji Kurisu, Michael Hippler, Jan Michael Schuller

**Affiliations:** Institution of Plant Biology and Biotechnology, University of Muenster, Muenster, Germany; Center for Synthetic Microbiology (SYNMIKRO) and Department of Chemistry, Philipps-University Marburg, Marburg, Germany; Institute for Protein Research, Osaka University, Suita, Osaka, Japan; Instisute of Plant Science and Resources, Okayama University, Okayama, Japan

**Author notes:** These authors contributed equally to this work (Yu Ogawa, Gyana Prakash Mahapatra, Yuval Milrad).

## Abstract

Photosynthetic electron transfer relies on small soluble carriers that shuttle electrons between the cytochrome *b*₆*f* complex and Photosystem I (PSI). While copper-containing plastocyanin (Pc) serves this role in plants, algae and cyanobacteria employ the heme protein cytochrome *c*₆ (Cyt *c*₆) as well. Here we present a cryo–electron microscopy structure of a Cyt *c*₆:PSI complex from *Chlamydomonas reinhardtii*. The Cyt *c*₆ heme is positioned ∼11 Å from P700, stabilized by extensive contacts involving the N-terminal domain of PSAF. Importantly, R66 in Cyt *c*₆, a key residue in ancestral donors, forms a putative electrostatic contact with PsaB-D623 and participates in a tri-planar π-stacking interaction with nearby aromatic residues. Our findings provide a structural framework for ancestral PSI interactions and illuminate the evolutionary diversification of electron transfer pathways.

## Main Text

The photosynthetic electron transport chain proceeds through a series of finely tuned bimolecular electron transfer reactions essential for maintaining efficient photosynthesis. This chain begins with Photosystem II (PSII), which catalyzes the oxidation of water and the reduction of plastoquinone (PQ). The reduced plastoquinol (PQH₂) is then oxidized by the Cytochrome *b*₆*f* complex (Cyt *b*₆*f*), transferring electrons to water-soluble carriers such as cytochrome *c*_6_ (Cyt *c*_6_) or plastocyanin (Pc), which in turn donate electrons to Photosystem I (PSI). This electron transfer through Cyt *b*₆*f* is coupled to proton translocation across the thylakoid membrane, generating an electrochemical proton gradient that drives synthesis of adenosine triphosphate (ATP).

From an evolutionary standpoint, the ancestral PSI is thought to have interacted with the heme-based electron carrier Cyt *c*₆(*1, 2*). However, over time due to selective pressures such as fluctuations in metal bioavailability, the copper-containing protein Pc emerged as a functional replacement for Cyt *c*₆. In contemporary photosynthetic organisms, this transition is reflected in the diversity of electron carriers; certain cyanobacteria still use only Cyt *c*_6_, other cyanobacterial and algal species switch between Cyt *c*_6_ and Pc depending on iron/copper availability, while vascular plants rely exclusively on Pc(*3*). Despite differences in metal cofactors, Cyt *c*_6_ and Pc have undergone convergent evolution and exhibit similar reaction kinetics with their partners, although these kinetics vary significantly among species(*1, 2, 4, 5*). In particular, PSI reduction by Cyt *c*_6_/Pc has been extensively studied using laser-flash absorption spectroscopy. In green algae and vascular plants, the kinetics typically show a fast first-order intramolecular electron transfer followed by the slower second-order bimolecular reaction(*2, 4–6*). The former suggests stable donor:PSI complex formation, while the latter reflects the interaction of freely diffusing donors and PSI. In contrast, in many cyanobacteria, the slower biomolecular phase is dominant or exclusive particularly *in vitro*(*2, 4, 5*).

From a structural perspective, deciphering the form of these donor:PSI complexes proved to be challenging. Early biophysical studies, accompanied by investigations of mutagenesis variants, concluded that the formation of these complexes is governed by a planar hydrophobic patch on the electron carriers (Cyt *c*_6_/Pc) docking into a hydrophobic groove at the PsaA–PsaB interface(*7–14*). Moreover, in the case of eukaryotic systems, the interaction is dominated by negatively charged residues, which form salt bridges with Lys residues in the extended N-terminal region of PSAF(*9, 11, 12, 15–20*). Recently, these estimations gained validation, as the structures of eukaryotic Pc:PSI complexes from both *Pisum sativum* and *Chlamydomonas reinhardtii* were resolved via cryo-EM (Cryogenic electron microscopy)(*21–23*). However, to date, there is no such structural evidence, showing exactly how Cyt *c*₆ bounds to PSI, as most recent attempts (mainly on cyanobacterial systems) have yet to meet a resolution, that would be sufficient for drawing biophysical conclusions(*24, 25*). This gap in structural knowledge still limits our understanding of the evolution and mechanistic diversity of donor:PSI reactions across photosynthetic lineages.

To address this gap, we performed cryo-EM analysis of a *Chlamydomonas reinhardtii* Cyt *c*₆:PSI complex, stabilized via chemical crosslinking that preserves a conformation competent for efficient electron transfer(*18, 26*). This work reveals that in *Chlamydomonas*, Cyt *c*_6_ utilizes the negatively charged residues interacting with PSAF similarly to eukaryotic Pc, while it retains an Arg residue that also significantly contributes to binding and electron transfer to PSI. The Arg is well conserved and crucial for the electron transfer in cyanobacterial donors but absent in eukaryotic Pc (Fig. S1)(*13, 27, 28*). Therefore, our structure offers a unique structural window into ancestral donor:PSI interactions, providing a structurally tractable model of their evolution.

## Results

### *Chlamydomonas* Cyt *c*_6_: PSI complex

To obtain structural insights into the Cyt *c*₆:PSI interactions, we reconstituted a *Chlamydomonas* Cyt *c*₆:PSI complex by chemical crosslinking, preserving a conformation allowing rapid electron transfer(*18, 26*). PSI (PsaB-His_20_) was affinity-purified from *Chlamydomonas* cells grown under standard conditions, while *Chlamydomonas* Cyt *c*_6_ was recombinantly expressed, purified, and pre-activated with EDC and sulfo-NHS(*23*). After removing excess crosslinking agents, the activated Cyt *c*_6_ was incubated with isolated PSI complex, enabling amide bond formation between Cyt *c*₆ carboxyl groups and PSI amino groups(*17, 18, 26*). The cross-linked Cyt *c*_6_:PSI complex was subjected through sucrose density gradient (SDG) centrifugation to separate it from the non-cross-linked Cyt *c*_6_, with the major PSI fraction recovered at lower density (Fig. 1A). Successful crosslinking was confirmed by the appearance of a ∼28.5 kDa band corresponding to the Cyt *c*₆:PSAF product and by the concurrent depletion of the free PSAF band (∼20 kDa) (Fig. 1B)(*18*). Thereafter, the cross-linked complex was subsequently subjected to cryo-EM single particle analysis (SPA) for structural characterization (Fig. S2).

**Fig. 1.**
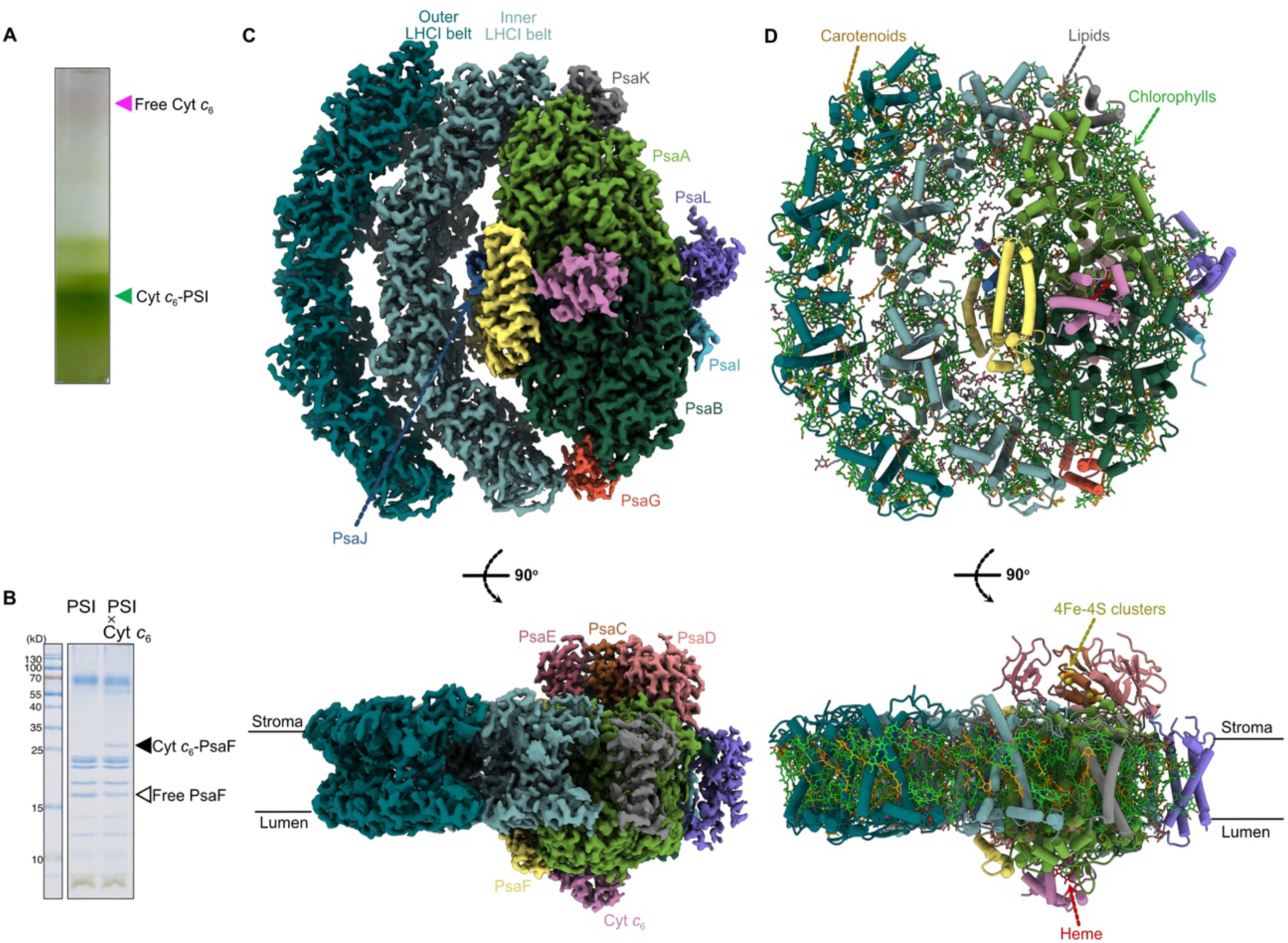
Purification and overall structure of Cyt *c*_6_:PSI complex. (**A**) Purification of cross-linked Cyt *c*_6_:PSI particles by sucrose density gradient centrifugation. (**B**) SDS-PAGE of PSI and cross-liked Cyt *c*_6_:PSI particles. (**C**) Electron density map of Cyt *c*_6_:PSI complex (top: luminal view; bottom: side view). (**D**) Structural model of Cyt *c*_6_:PSI complex (top: luminal view; bottom: side view).

Cryo-EM SPA of the sample yielded a reconstructed map with a global resolution of 1.83Å, in which PSI was resolved with near-atomic detail (Fig. 1, C and D; Fig. S2-4). The structure closely resembled previously reported *Chlamydomonas* PSI monomers, and the map quality allowed for precise modeling of all core subunits and cofactors (Fig. 1, C and D; Fig. S4)(*23, 29–32*). The PSI model comprises 19 protein subunits and includes 189 chlorophyll a, 112 chlorophyll b, 27 β-carotenes, 2 neoxanthins, 7 xanthophylls, 16 luteins, 2 phylloquinones, 3 [4Fe–4S] clusters, and 63 lipid molecules, along with 1,088 water molecules. Although weak density was initially observed near PSAF in the global map, consistent with Cyt *c*₆, the signal was insufficient for confident interpretation (Fig. S2). To improve visualization of Cyt *c*₆, we performed focused refinement in CryoSPARC using a soft mask around the expected binding site(*33, 34*). This yielded a locally enhanced reconstruction at 2.06 Å, in which an additional density adjacent to PSAF was clearly resolved (Fig. S3). The shape and volume of the density matched that of Cyt *c*₆, enabling us to rebuild the Cyt *c*₆ model into the density, based on the *Chlamydomonas* Cyt *c*₆ crystal structure(*35*).

In addition to the luminal-side Cyt *c*₆ density, we also observed two non-protein assigned densities on the stromal side of the PSI complex, located near the PSAD and PSAE subunits (Fig. S2C). These densities were consistently present across multiple reconstructions but remained insufficiently resolved for interpretation, likely due to extreme conformational flexibility or heterogeneous occupancy. Their position is reminiscent of previously reported stromal-side densities in PSI structures from *Chlamydomonas*, where they were proposed to correspond to transient or regulatory interaction partners such as FNR or loosely bound ferredoxin(*36*). While the exact identity of the stromal densities in our structure remains unclear, their consistent localization suggests a potential functional relevance.

### Cyt *c*_6_:PSI binding

The donor binding site of PSI accommodates a shallow hydrophobic pocket, formed at the luminal interfaces of PsaA and PsaB subunits (Fig. S5A, gray circle). This pocket is delineated at its base by the symmetrical *l*/*l’* loops (Fig. S5B, gray and black brackets, respectively), which hold a Trp dimer in its center (PsaA-W651 and PsaB-W626, Fig. 2A; Fig. S5B). This region is enclosed by positively-charged protrusion of the N-terminal domain of PSAF. It is important to note that the PsaA-PsaB region is highly conserved throughout the entire photosynthetic domain, and that the bulky edge of PSAF is present in vascular plants and algae (Fig. S1)(*37*). Our data show that the binding of Cyt *c*_6_ to PSI is stabilized by an extensive network of non-covalent interactions. Our high-resolution density map show a well-defined density corresponding to the covalent cross-linking between PSAF-K27 and Cyt *c*₆-E69, which also interacts with PSAF-K23, though the crosslinking was not visually detectable (Fig. 2B, note that here for consistency with earlier reports, we count PSAF residues starting from the N-terminus of the mature protein eliminating the transit peptide segment, which is included in the PDB data and analysis). The involvement of other specific residues and water molecules was identified using LigPlot^+^ software (Fig. S6 and S7A)(*38, 39*). Interestingly, other charged residues on PSAF and Cyt *c*₆, were not located in sufficient proximity that enables salt bridge formation (<6Å) (Fig. 2B)(*40, 41*). However, this observation should be addressed with caution, as the experimental procedure might have had an effect in this respect.

**Fig. 2.**
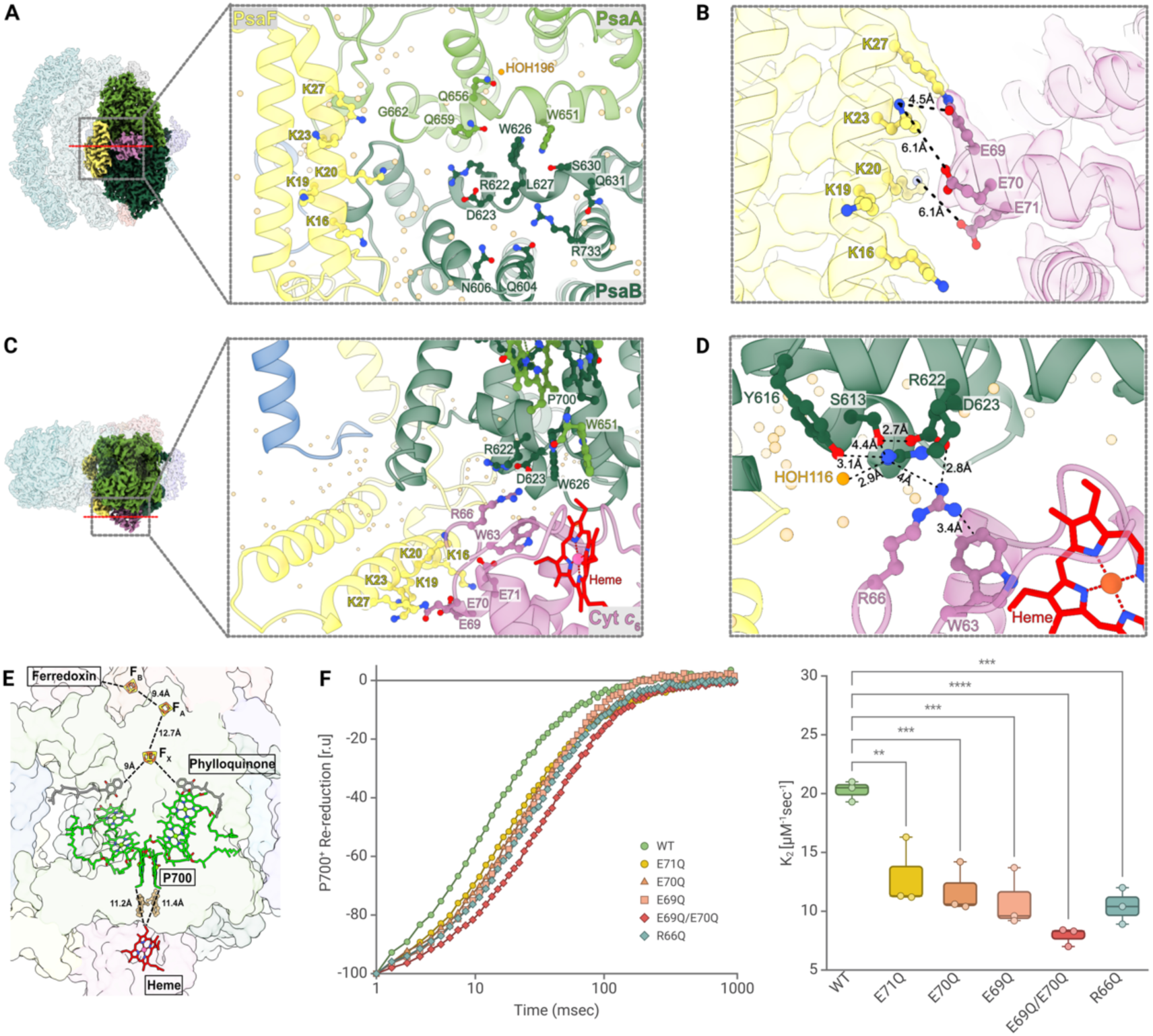
Cyt *c*_6_:PSI binding. (**A**) Luminal view of Cyt *c*_6_:PSI interface (just PSI side). Directly interacting amino acids and water molecules (orange) are shown, in addition to PsaA-W651 and PsaF-K16/K19/K20. (**B**) Cyt *c*_6_:PsaF interface with the corresponding cryo-EM density. The putatively involved residues are shown with their distances. (**C**) Side view of Cyt *c*_6_:PSI interface. (**D**) Cyt *c*_6_:PsaB interface. The amino acid residues and a water molecule putatively involved in the protein-protein interactions mediated by R66 are shown with their distances. (**E**) Edge-to-edge distances of electron-transfer molecules in Cyt *c*_6_:PSI complex. (**F**) *In vitro* kinetic analysis of P700^+^ re-reduction using Cyt *c*_6_ variants with mutations at putatively interacting amino acid residues. Box-plots show the average second-order rate constants (k_2_). Statistical analysis was conducted using a One-way ANOVA with Dunnett multiple comparisons test. The illustration and statistical analysis were generated using BioRender.com.

Within the shallow pocket, on the PsaA-*l* loop, a water-mediated hydrogen bond (HOH196) connects the PsaA-Q656 and Cyt *c*₆-G58, while PsaA-Q659 hydrophobically interacts with Cyt *c*₆-P61/A62 (Fig. S6A and S7B, left). The PsaB-*l’* loop provides more extensive contacts among non-charged residues and the heme group of Cyt *c*₆ (Fig. S6, B and D and S7B, right). One interesting exception is the case of a conserved charged pair of PsaB-R622/D623, that interacts with Cyt *c*₆-R66; although PsaA-*l* loop also contains the structurally symmetric Arg/Asp pair (R647/D648), it is not directly engaged in Cyt *c*₆ binding. The interface over the shallow pocket is further reinforced by additional peripheral contacts; specifically, PsaA-G662 interacts with Cyt *c*₆-D65, PsaB-Q604/606 with Cyt *c*₆-N11/G12, and PsaB-R733 with Cyt *c*₆-A16 (Fig. S6, A and B and S7C). Cyt *c*₆-D65 also forms a hydrogen bond with PSAF-K20, extending the peripheral contact site on PsaA into PSAF (Fig. S6C and S7C, top).

The aforementioned R622/D623 pair on the PsaB-*l’* loop engages Cyt *c*₆-R66, which was shown to be critical for PSI binding and electron transfer in cyanobacteria (Fig. 2, C and D; Fig. S6B)(*27, 28*). To better understand the functional importance of this residue, we examined its local environment beyond the direct PsaB interface (Fig. 2D). Cyt *c*₆-R66 may form an electrostatic interaction with PsaB-D623 while also contacting PsaB-R622. These two residues could partially neutralize each other. Additionally, the hydroxyl groups of PsaB-S613 and PsaB-Y616 as well as a water molecule (HOH116) can further delocalize the local charges. This microenvironment suggests that electrostatics alone may not fully account for the essential role of the corresponding Arg residue in cyanobacteria. Notably, within Cyt *c*₆ itself, W63 lies directly beneath R66 and may participate in π(cation)-π interactions(*42, 43*), and in the Cyt *c*₆:PSI complex, the indole group of W63 and the guanidinium groups of R66 and PsaB-R622 are arranged in a semi-parallel stack at distances of 3–5 Å, suggesting a potential three-layer π(cation)-π interactions that could further stabilize the donor:acceptor interface. While PsaB-R622 was previously predicted to interact with Cyt *c*₆-D65(*13*), this residue instead makes the peripheral contacts on PsaA and PSAF as mentioned above (Fig. S6, A and C, and S7C, top).

Together, the Cyt *c*₆:PSI binding is supported by a broad, multivalent interaction surface encompassing polar, charged, hydrophobic and solvent-mediated elements, as well as π(cation)-π interactions. In the complex, the redox cofactors which are involved in interprotein electron transfer (i.e. the heme of Cyt *c*₆ and the P700 chlorophyll pair of PSI) are positioned at an edge-to-edge distance of 11.2-11.4 Å, placing them in a suitable range for efficient electron transfer (Fig. 2E). We also observed that these two cofactors encompass the Trp dimer situated in the center of the *l*/*l^’^* loops. Moreover, the tight contact on this region excludes surrounding water molecules from the redox interface, presumably lowering the energetic barrier for electron transfer (Fig. 2A; Fig. S5B).

### Functional significance of eukaryote/cyanobacteria-like interactions

Green algal Cyt *c*₆ is uniquely positioned in evolution, utilizing both PSAF-mediated electrostatic interactions, while simultaneously conserving R66, a residue known to be essential for PSI interaction in cyanobacteria (Fig. S1)(*13, 26*). To test the role of these interactions, we generated several single mutations on Cyt *c*₆, which eliminate its negative charge: E69Q, E70Q, E71Q, in addition to a double mutant: E69Q/E70Q. Moreover, to verify the role of R66, we generated another mutant variant, namely R66Q. To evaluate the impact of these mutations, we measured P700^+^ re-reduction kinetics (Fig. 2F) and assessed second-order rate constants (k₂)(*37*). Our results show that all variants exhibit a significant decrease of k₂ compared to wild-type Cyt *c*₆, with the most severe impairment observed for the E69Q/E70Q double mutant. The E71Q variant showed a more modest reduction, suggesting a less critical role for this residue.

We then assessed the impact of these mutations on complex stability using chemical cross-linking (Fig. S8). Accordingly, R66Q, E69Q, and E70Q mutations drastically affected the cross-link formation between Cyt *c*₆ and PSAF, supporting their direct involvement in intermolecular interactions. In contrast, E71Q showed little effect on cross-linking efficiency, consistent with its weaker kinetic phenotype. Interestingly, the phenotype of E69Q/E70Q double mutant was weaker than those of each single mutant, suggesting that compensatory interactions or altered residue orientation may permit residual complex formation.

Overall, these results demonstrate that the interactions mediated by the conserved negative residues (in particular, E69 and E70) and R66 of Cyt *c*₆ are both critical for stable PSI binding and efficient electron transfer. These observations underscore green algal Cyt *c*₆ as a unique evolutionary intermediate of PSI donors, which harnesses both the novel and ancestral features.

## Discussion

In this work, we present a high resolution cryo-EM structures of the Chlamydomonas PSI core at 1.83 Å and its Cyt *c*₆-bound complex at 2.06 Å, revealing detailed molecular interactions where the novel and ancestral features coexist. This high resolution enables us to draw new conclusions about donor:PSI complex formation. Of note, the conformation of Cyt *c*₆ binding reported here is dissimilar to those predicted by modeling in previous studies (*44–46*). Here, we observed that the interface is extensively stabilized by a network of hydrogen bonds, salt bridges, hydrophobic contacts and π(cation)-π interactions primarily involving the *l*/*l’* loops and the N-terminal domain of PSAF (Fig. 2; Fig. S5-7). The comprehensive disclosure of the interactions within the shallow pocket indicates on an asymmetric contribution from PsaA and PsaB: as the latter subunit provides a greater number of interacting residues (Fig. 2A, S6 and S7). Cyt *c*₆ adheres more closely to PsaB and contacts with PsaA at limited sites, while positioning the heme right beneath the Trp dimer and P700 (Fig. 2E). This asymmetry may explain the stronger functional impact of PsaA-W651 mutations on Cyt *c*₆ binding and electron transfer relative to PsaB-W626(*10, 13*). This interaction network ensures the stringent donor:acceptor orientation; the Cyt *c*₆ heme is positioned approximately 11 Å from the P700 chlorophyll dimer, with the conserved Trp residues located along the electron transfer axis. This architecture is consistent with reports of rapid electron transfer kinetics (∼4 μs), a characteristic of native as well as of cross-linked first-order Cyt *c*₆:PSI complex(*18, 26*).

Superimposition of Cyt *c*₆:PSI on the Pc:PSI from Chlamydomonas (PDB: 7ZQE, 7ZQC) enabled us to compare the electron transfer pathways created by the alternative donors (Fig. 3)(*21*). First, we observed the co-localization of the heme iron of Cyt *c*_6_ and the copper of Pc, positioned directly below the Trp gateway and P700. In addition, we noted similar exclusion of surrounding water molecules from the interface, which couples electron transfer to minimal solvent reorganization (Fig. 2A; Fig. S5B)(*22, 23*). Taken together, these observations suggest that transferred electrons might be relayed *via* π-conjugated indole system *en route* to the reaction center, highlighting the functional equivalence of these two evolutionary distinct donors.

**Fig. 3.**
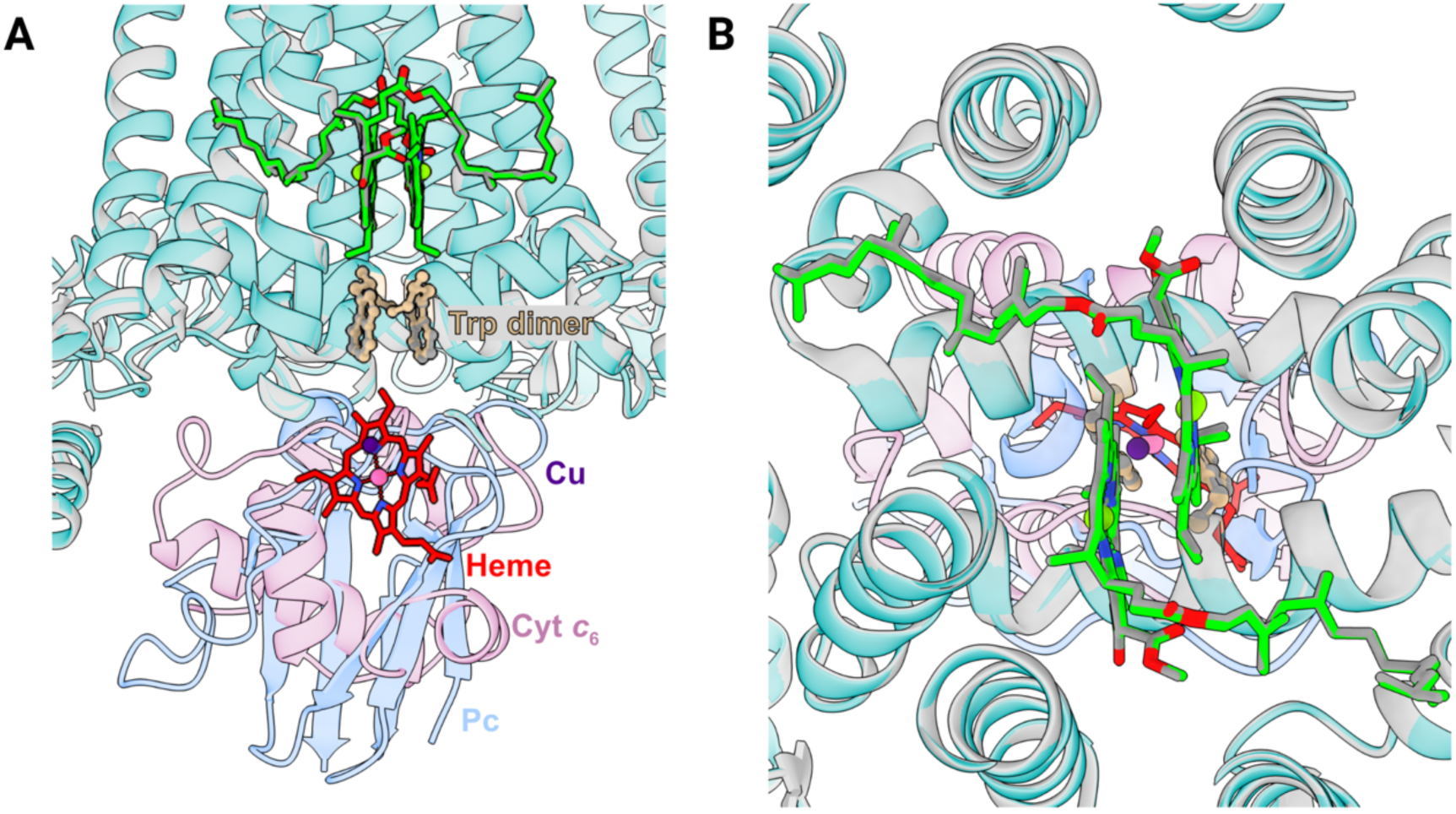
Comparative overlay of donor:PSI complexes. (**A**) Overlay of the Cyt *c*_6_:PSI on the *Chlamydomonas* Pc:PSI (PDB ID 7ZQE, 7ZQC). The former PSI is colored in light blue, the latter in gray. (**B**) Stromal view of the overlayed structures of *Chlamydomonas* Cyt *c*_6_:PSI and Pc:PSI.

The structure also allows dissecting the electrostatic interactions mediated by the N-terminal domain of PSAF, a characteristic of vascular plants and green algae. The covalent cross-link between PSAF-K27 and Cyt *c*₆-E69, resolved by cryo-EM captures a dominant and highly uniform binding conformation, validating the complex structure as a faithful representation of the physiological electron transfer assembly (Fig. 2B). Mutagenesis analysis revealed that E69 and E70 are critical for complex formation and electron transfer, while E71 plays a more limited role (Fig. 2F; Fig. S8). Additionally, positively charged residues on the N-terminal domain of PSAF were previously shown to be necessary for complex stability and efficient electron transfer(*19*). However, despite their important role, charged residues other than PSAF-K23/K27 and Cyt *c*₆-E69 were too far apart to form salt bridges (Fig. 2B). Possibly, they may be required only to form a transient and loose contact between donor and acceptor prior to the final conformation, which is competent for electron transfer. A similar conclusion was drawn in a previous NMR study on plant Pc as well(*47*). The unexpected cross-link formation by the E69Q/E70Q Cyt *c*₆ variant also suggests other complex conformations, which needs architectural change for electron transfer (Fig. S8).

Our structural data further provide the first direct visualization of Cyt *c*₆-R66-mediated interactions, previously implicated in cyanobacterial systems, revealing a putative electrostatic contact with PsaB-D623 (Fig. 2D; Fig. S6B), a residue whose involvement had been proposed but not structurally confirmed(*13*). At the same time, R66 can engage in a three-layer π(cation)-π interaction along with PsaB-R622 and Cyt *c*₆-W63 (Fig. 2D; Fig.S6B). The dual strong interactions mediated by R66 may explain its crucial role. While R66 appears less critical in Chlamydomonas than its cyanobacterial counterparts, where substitution of the analogous Arg residues in *Synechocystis* sp. PCC 6803 and *Anabaena* sp. PCC 7119 leads to marked diminishments in electron transfer rates(*27, 28*), our data indicate that R66 nonetheless contributes to PSI association and facilitates electron transfer in the eukaryotic complex (Fig. 2F; Fig. S8). Moreover, cyanobacterial Pc also contains the crucial Arg residue (R88) corresponding to Cyt *c*₆-R66, along with the underlying aromatic residue (Y83) equivalent to Cyt *c*₆-W63(*42*). This suggests that a similar three-layer π(cation)-π interaction is possible for cyanobacterial Pc. Notable is that the π interactions exist nearby the redox cofactor, heme or copper; the subjacent aromatic residue (Pc-Y83/Cyt c_6_-W63) is an immediate neighbor of the cofactor (Fig. 2, C and D)(*42*). In this regard, it was reported that the redox state of cyanobacterial Pc changes the conformation of R88, suggesting the electron transfer reaction can modify the π(cation)-π interactions (*48*). Furthermore, similar series of π(cation)-π interactions within the vicinity of electron transfer pathways is also implicated in ferredoxin–PSI interface(*24*). Therefore, it is tempting to hypothesize that π electrons act as key modulators, coordinating donor:acceptor binding and electron transfer reactions. This could explain how reduced donors efficiently bind to acceptors, while their oxidized form unbinds swiftly, optimizing the turn-over of these reactions (*6, 49*). However, the functional significance of the π interactions is partly buried by the PSAF-mediated interactions in Chlamydomonas Cyt *c*_6_ (Fig. 2F; Fig. S8), and in eukaryotic Pc, the Arg residue is lost and completely replaced by the long-range electrostatic interaction, which is more powerful considering that it is attributed to the stable donor:PSI complex formation observed in vascular plants and green algae (Fig. S1)(*20, 50*).

Together, these findings illuminate an evolutionary trajectory wherein the emergence of PSAF-dependent interactions enhanced both binding affinity and electron transfer efficiency, reducing the required concentration of electron donors and economizing iron and copper use in chloroplasts relative to cyanobacterial cells.

## Acknowledgments

We thank Christoph Gerle for constructive and insightful discussions, Laura Mosebach and Vida Adlfar for experimental supervision, and Rémi Ruedas and Mohamed Chami for collecting the cryo-EM data at the BioEM lab of the Biozentrum, University of Basel.

## Author contributions

Conceptualization: MH

Methodology: GPM, JMS, GK

Investigation: YO, GPM, YM, MS

Visualization: YO, GPM, YM

Funding acquisition: MH, JMS

Project administration: MH, JMS

Supervision: MH, JMS

Writing – original draft: YO, GPM, YM

Writing – review & editing: MH, JMS

## Data and materials availability

The raw data supporting this article will be made available by the authors, without undue reservation.

## Supplementary Materials

Materials and Methods

Supplementary Text

Figs. S1 to S8

Tables S1

References (*51–61*)

Data S1

## Materials and Methods

### PSI purification

Experiments were performed using the previously mentioned *Chlamydomonas* strain that expresses PsaB fused with His_20_-Tag after the third residue from the N-terminus(*23*). They were kept on Tris-acetate-phosphate (TAP) medium, solidified with 1.5% w/v agar at 25 °C under light of ∼20 μmol photons m^-2^ s^-1^. For experiments, the algae were cultured in liquid TAP medium on a rotary shaker (120 rpm) at 25 °C under continuous light of ∼20 μmol photons m^-2^ s^-1^.

The PSI purification was performed according to the protocol mentioned previously with some modifications(*23*). The *Chlamydomonas* cells were harvested under oxic conditions by centrifugation at 3,000 rpm at 4°C for 5 min (Beckman Coulter JLA-16.250 rotor). They were resuspended in ice-cold buffer (25 mM HEPES/KOH, pH 7.5, 0.33 M sorbitol, 5 mM MgCl_2_, 1 mM PMSF, 1mM benzamidine and 5 mM aminocaproic acid) and homogenized using a nebulizer (2 bar, two passages). The nebulized cells were collected by centrifugation at 20,000 rpm at 4°C for 10 min (Beckman Coulter JA-25.50 rotor), and the pellets were thoroughly resuspended using a potter homogenizer in 5 mM HEPES/KOH, pH 7.5, 0.5 M sorbitol, 10 mM EDTA, 1mM benzamidine and 5 mM aminocaproic acid). The resuspended material was layered on top of a sucrose density gradient (1.8 M and 1.3M sucrose, the same composition as the resuspension buffer). After ultracentrifugation at 24,000 rpm at 4 °C for 1 h (Beckman Coulter SW 32 Ti rotor), the thylakoid membranes were recovered by pipetting the step gradient interphases and diluted ∼4 times, followed by pelleting by centrifugation at 21,500 rpm at 4 °C for 20 min. The isolated thylakoids were carefully resuspended using a paint brush and set to 1 mg Chl mL^-1^ in 5 mM HEPES/KOH, pH 7.5, 1mM benzamidine and 5 mM aminocaproic acid. The chlorophyll concentration was determined using spectrophotometer(*51*). The same volume of 2% [w/v] α-DDM in a buffer of the same composition was added, and the thylakoids were solubilized on ice for 10 min with occasional gentle mixing. Un-solubilized materials were precipitated by centrifuging at 14,000 rpm at 4 °C for 5 min, and the supernatant were diluted 5 times in 25 mM HEPES/KOH, pH 7.5, 100 mM NaCl, 5 mM MgSO_4_, 10% [v/v] glycerol, 1mM benzamidine and 5 mM aminocaproic acid. The sample was loaded onto a TALON metal affinity column (1 ml resin mg Chl^-1^) at a rate of ∼0.5 ml min^-1^. The column was washed with ten times the volume of 25 mM HEPES/KOH, pH 7.5, 100 mM NaCl, 5 mM MgSO_4_, 10% [v/v] glycerol and 0.02% [w/v] α-DDM at a rate of ∼0.5 ml min^-1^ and then washed with the same volume of 25 mM HEPES/KOH, pH 7.5, 100 mM NaCl, 5 mM MgSO_4_, 10% [v/v] glycerol, 0.02% [w/v] α-DDM and 5 mM imidazole at a rate of ∼1 ml min^-1^. The PSI was eluted with 25 mM HEPES/KOH, pH 7.5, 100 mM NaCl, 5 mM MgSO_4_, 10% [v/v] glycerol, 0.02% [w/v] α-DDM and 150 mM imidazole. The PSI was concentrated with a spin column (regenerated cellulose: 100,000 molecular weight cut-off (MWCO)). The sample was diluted ∼10 times with 30 mM HEPES/KOH, pH 7.5, 0.02% [w/v] α-DDM and reconcentrated twice.

### Cyt *c*_6_ mutation and purification

To heterologously express *Chlamydomonas* Cyt *c*_6_, the wild type (WT) Cyt *c*_6_ CDS was synthesized (GenScript) and ligated into the NdeI/EcoRI restriction site of the expression vector pET22b. The resulting plasmids was used to transform NEB 5α competent *E. coli* cells. Site-directed mutagenesis for different Cyt *c*_6_ variants was performed employing In-Fusion Snap Assembly Starter Bundle (Takara). The pET22b-Cyt *c*_6_ (WT) was amplified using specific pairs of primers carrying the new codons for each Cyt *c*_6_ variants listed in Table S1. The linear amplicons were assembled into circular plasmids following the manufacturer’s instructions, to produce pET22b-Cyt *c*_6_ (R66Q), pET22b-Cyt *c*_6_ (E69Q), pET22b-Cyt *c*_6_ (E70Q), pET22b-Cyt *c*_6_ (E71Q) and pET22b-Cyt *c*_6_ (E69Q/E70Q), which were used to transform Steller Competent Cells (Takara). The plasmids were isolated from the competent cells and confirmed by sequencing (Eurofins Genomics). For recombinant protein expression, NEB BL21(DE2) competent *E. coli* cells were co-transformed with each of the pET22b plasmids and pEC86, which expresses *E. coli* ccm genes and improve expression of mature c-type cytochromes in the same organism(*52*). The protein expression was induced by culturing the *E. coli* strains in LB medium containing 1 mM IPTG, 50 μgr mL^-1^ Ampicirin and 34 μgr mL^-1^ Chloramphenicol on a rotary shaker (140 rpm) at 30 °C overnight. The cells were harvested by centrifugation at 5,000 rpm at 4 °C for 5 min (Beckman Coulter JLA-16.250 rotor). The pelleted cells were resuspended in lysis buffer (20 mM Tricine, pH 7.8, 10 mM KCl and 1 mM PMSF) and sonicated for a total of 180 s (Brason 250 Digital Sonifier w/ Prob, Marshall Scientific). The homogenate was centrifuged at 10,000 rpm at 4 °C for 30 min (Beckman Coulter JA-25.50 rotor), and the supernatant was loaded on an anion exchange column (DEAE Sepharose CL-6B, GE Healthcare) at a rate of ∼1 ml min^-1^. The column was washed with ten times the volume of the lysis buffer, and the second wash was done with 20 mM Tricine, pH 7.8 and 50 mM KCl. The Cyt *c*_6_ was eluted with 20 mM Tricine, pH 7.8 and 400 mM KCl and was concentrated using a spin column (regenerated cellulose: 3,000 MWCO). The sample was diluted ∼10 times with 20 mM Tricine, pH 7.8 and 10 mM KCl and reconcentrated. The sample was further purified employing a size exclusion chromatography (Superdex 65 10/300 GL on an AKTA pure system). The concentration of Cyt *c*_6_ was determined spectroscopically (at 552 nm with an extinction coefficient of 20 mM^-1^ cm^-1^)(*53*).

### Cross-linking and Cryo-EM sample preparation

Cyt *c*_6_:PSI cross-linking was performed as previously described with some modifications(*23*). The *Chlamydomonas* WT Cyt *c*_6_ was loaded on a PD G25 desalting column and eluted with 30 mM HEPES/KOH, pH 7.5, followed by concentration using a spin column (regenerated cellulose: 10,000 MWCO). The Cyt *c*_6_ (100 μM) was preactivated by incubating with 5 mM 1-ethyl-3-[3-dimethylaminopropyl] carbodiimide (EDC) and 10 mM sulfo-N-hydroxysuccinimide (NHS) for 20 min at room temperature in dark. Un-reacted crosslinkers were removed using a PD G25 desalting column, and the activated Cyt *c*_6_ was concentrated using a spin column (regenerated cellulose: 10,000 MWCO). The activated Cyt *c*_6_ (20 μM) and the purified PSI (0.1 mg Chl mL^-1^) were mixed in 30 mM HEPES/KOH, pH 7.5 containing 1mM ascorbate, 0.1 mM DAD, 3 mM MgCl_2_ and 0.03% [w/v] α-DDM, and the mixture was incubated for 45 min at room temperature in dark for a cross-linking reaction, followed by purification via a 1.3 M to 0.1 M sucrose density gradient including 5 mM HEPES/KOH, pH 7.5 and 0.02% [w/v] α-DDM (∼60 mg Chl per gradient). The PSI fractions were collected after ultracentrifugation at 33,000 rpm at 4 °C for 14 h (Beckman Coulter SW 41 Ti rotor). For Cryo-EM analysis, the sample was subjected to PD G25 desalting column to remove sucrose and subsequently concentrated to ∼1.6 mg Chl mL^-1^ in 34 μl.

For the crosslinking experiments in Fig. S8, non-activated Cyt *c*_6_ (4 μM) and PSI (40 μg Chl mL^-1^) were incubated in 30 mM HEPES/KOH, pH 7.5 containing 1mM ascorbate, 0.1 mM DAD, 3 mM MgCl_2_, 0.03% [w/v] α-DDM, 5 mM 1-ethyl-3-[3-dimethylaminopropyl] carbodiimide (EDC) and 10 mM sulfo-N-hydroxysuccinimide (NHS) for 45 min at room temperature in dark. The cross-linking reaction was quenched with 50 mM Tris, pH 8.0, and the reaction mixture was subjected to SDS-PAGE and Western blotting.

### SDS-PAGE and Western blotting

The protein samples were supplemented with one tenth volume of loading buffer (250 mM Tris/HCl, pH 6.8, 8% [w/v] SDS, 40% [v/v] glycerol and 0.33% [w/v]) and 100 mM DTT and subsequently heated at 65 °C for 20 min. After separated in SDS-PAGE via electrophoresis, they were stained with Coomassie Serva Blue G or electro transferred onto nitrocellulose membranes (Amersham). For Western blotting, the membranes were incubated with specific antibodies, and the protein-antibody complexes were labeled using a Supersignal West Pico Plus chemiluminescent substrate (Thermo Scientific).

### Cryo-EM data acquisition, processing and model building

For cryo-EM sample preparation, 4 μl of purified Cyt *_C_*_6_:PSI complex was applied to glow-discharged Quantifoil R 2/1 copper 300-mesh holey carbon grids (PELCO easiGlow, 25 s at 15 mA). Grids were vitrified in liquid ethane-propane using a Vitrobot Mark IV (Thermo Fisher Scientific) at 100% humidity and 4 °C, with a blotting time of 9 s and blotting force 7. Cryo-EM data was collected on a Titan Krios G4 transmission electron microscope (Thermo Fisher Scientific) operated at 300 kV, equipped with a Selectris energy filter and Falcon 4 direct electron detector. Movies were acquired in counting mode at a nominal magnification of 165,000× yielding a calibrated pixel size of 0.73 Å. Each exposure was 1.9 s and split into 65 frames, with a total accumulated dose of 50 e−/Å². In total, 21,413 movies were collected.

All cryo-EM data processing was performed in CryoSPARC(*33*). Micrographs were gain-corrected, aligned, and dose-weighted using Patch Motion Correction(*54*). Thereafter, the contrast transfer function (CTF) was estimated using Patch CTF(*55*). After discarding micrographs with poor CTF fits or suboptimal quality, 21,147 micrographs were retained for particle picking. Initial particle picking was carried out on a subset of 1,000 micrographs using the Blob picker. 2D classes generated from this subset were used to train a Topaz model(*56, 57*), which underwent four rounds of iterative training. The final Topaz model was used to extract particles on the entire dataset, resulting in approximately 1.3 million particles. Particles were extracted with a 512-pixel box size, binned 4 times for initial classification. Extracted particles were subjected to 2D classification to remove contaminants. The retained particles after the 2D classifications were then subjected to five ab initio 3D reconstructions, resulting in both high-quality and junk classes. These reconstructions were refined through heterogeneous refinement. Multiple rounds of ab initio heterogeneous refinement were performed to systematically remove poor-quality classes. After these steps, around 246,000 high quality particles remained and were re-extracted at full (unbinned) 512-pixel box size for high-resolution refinement. These particles were refined by non-uniform refinement in CryoSPARC, yielding a reconstruction at 1.83 Å global resolution (PSI map), as estimated with the gold-standard Fourier shell correlation (FSC) with a cut-off of 0.143. At this point, the Cyt *c*_6_ component was not well resolved, therefore a mask was generated around Cyt *c*_6_, followed by masked 3D classification into six classes. In one class, Cyt *c*_6_ showed a clear well-resolved density; this class was further refined locally using the focused mask, giving a 2.06 Å map (Cyt *c*_6_-PSI-core map) where side chains, pigments, cofactors, and the crosslink between PsaF and Cyt *c*_6_ (Fig. S2) were well resolved. Two additional densities were observed on the stromal side of the complex. To further resolve these densities, focused classification was performed, however, due to extreme flexibility, they could not be refined. The resulting PSI and Cyt *c*_6_-PSI-core maps were used for model building.

For the PSI map, the previously determined *Chlamydomonas reinhardtii* PSI structure (PDB ID: 7ZQC) was used as the starting model(*23*). For the Cyt *c*_6_-PSI-core map, the same PSI structure was combined with the crystal structure of Cyt *c*_6_ (PDB ID: 1CYI)(*35*). Both models were rigid body fit into the 1.83 Å and 2.06 Å maps respectively using UCSF ChimeraX v1.10(*58*). Side chains and ligands were inspected and fitted manually in Coot v0.9.8.9(*59, 60*). Given the high resolution, water molecules were modeled automatically using the water picking tool in Coot, and each placement was inspected manually. Final models were refined using real-space refinement in PHENIX v1.21.1-4907(*61*). CIF restraints for chlorophyll B (CHL) and chlorophyll A isomer (CL0) were generated with the Grade Web Server (https://grade.globalphasing.org/), while those for lutein (LUT) and neoxanthin (NEX) were created using the eLBOW utility in PHENIX by providing the chemical file (mol2) and final geometry file from the PDB models as inputs. Before PDB submission, multiple rounds of validation and model building were carried out using Phenix and Coot. And finally, all figures were prepared using UCSF ChimeraX.

### Single-Flash Absorbance Spectroscopy

Kinetic studies of P700 post laser flash re-reduction we conducted as described previously(*37*). In short, isolated PSI complexes (331 nM) were solubilized in 7.5 mM KCl, 2.5 mM MgCl_2_, 10 mM ascorbate, 0.5 mM DAD and 1 mM methyl viologen, pH 7.0 (MOPS 5 mM). The mixture was placed in a “Joliot type Spectrophotometer” (JTS-150, Biologics) supplied with a “Smart-Lamp” with a dual measuring light usage (705 to 740 nm) and adequate detector filters (P700). Absorbance was measured post a laser flash (100 detections in a decreased exponential rate for the duration of 5 seconds with an initial delay of 700 µsec, 4 technical repetitions per test). Cyt *c*_6_ was gradually added (0.5-4.0 µM, Data S1) and k_2_ values were calculated using Origin.Pro (ExpDec1). For generating the boxplots, the k_2_ values for 2-4 µM Cyt *c*_6_ were averaged, using 3 biological replicates of independent PSI isolations. For statistical comparisons, a One-way ANOVA with Dunnett multiple comparisons test was applied using Biorender.com.

---

**Fig. S1.**
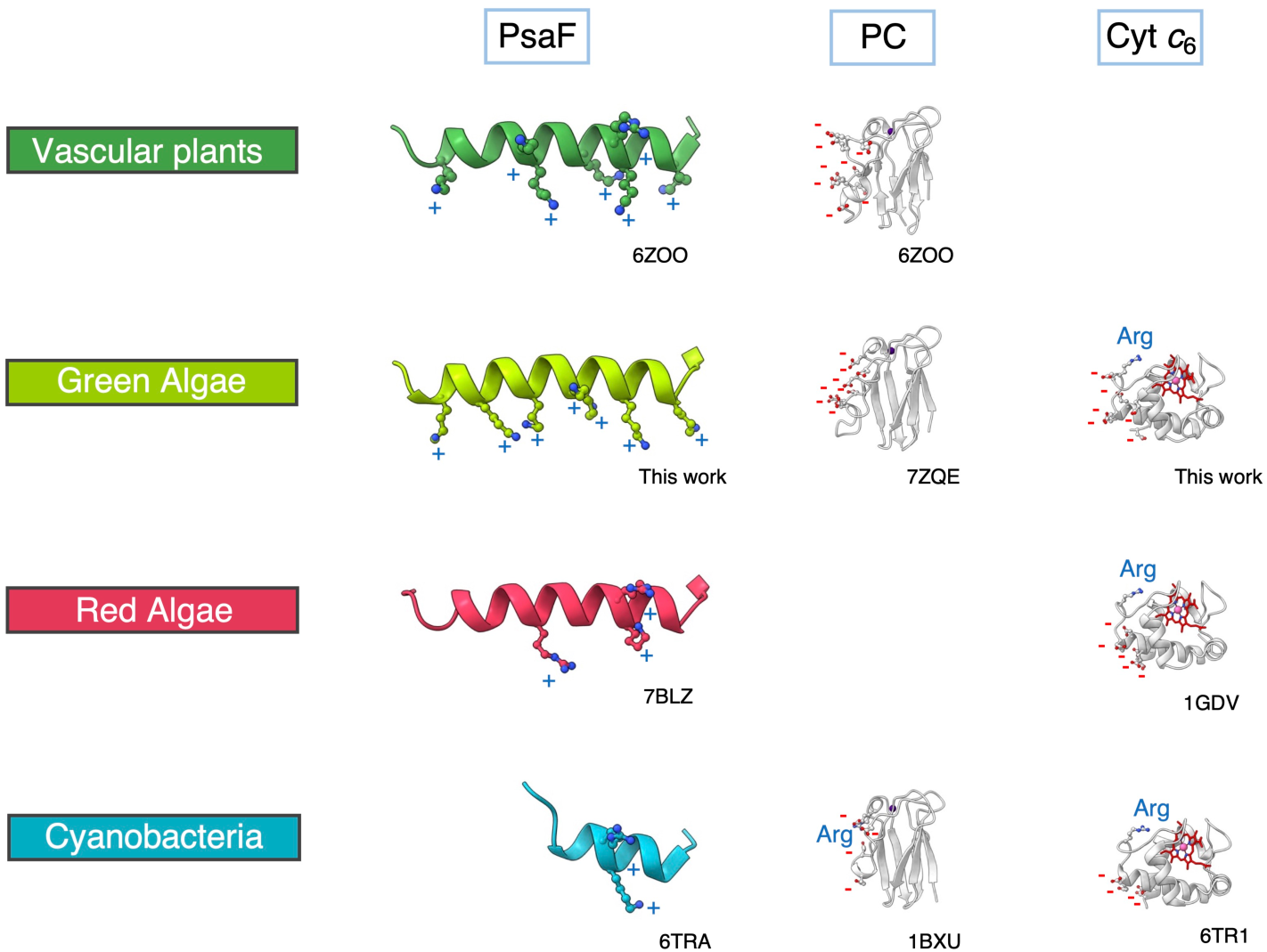
Comparison of the charged residues on interfaces of PsaF, PC and Cyt *c*_6_ from vascular plants, green algae, red algae and cyanobacteria. Each PDB ID is provided.

**Fig. S2.**
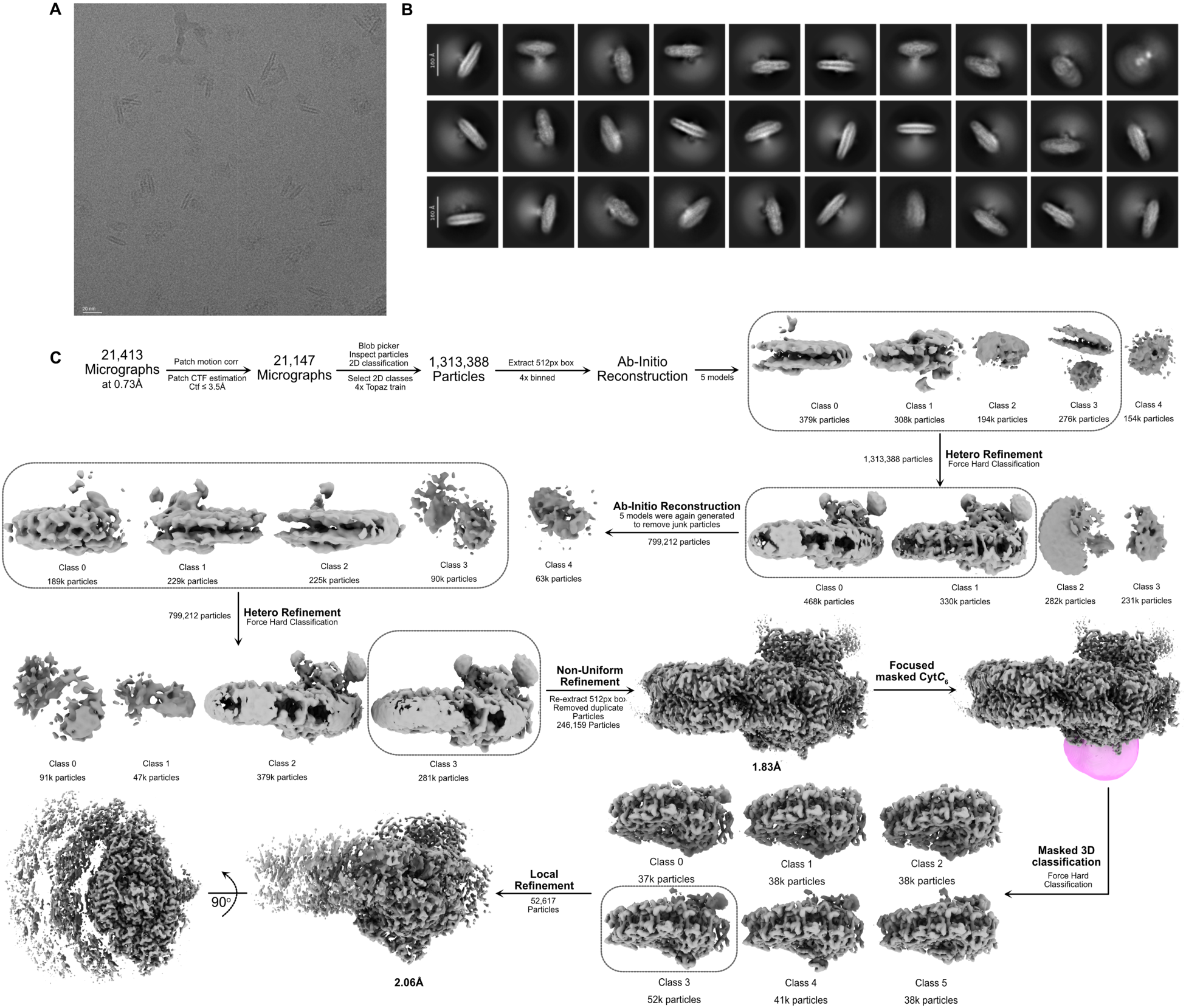
Cryo-EM data collection and processing workflow. (A) A representative micrograph showing Cyt *c*_6_:PSI particles. (B) Reference-free 2D class averages revealing different views of the complex. (C) Overview of the cryo-EM data processing pipeline performed in cryoSPARC.

**Fig. S3.**
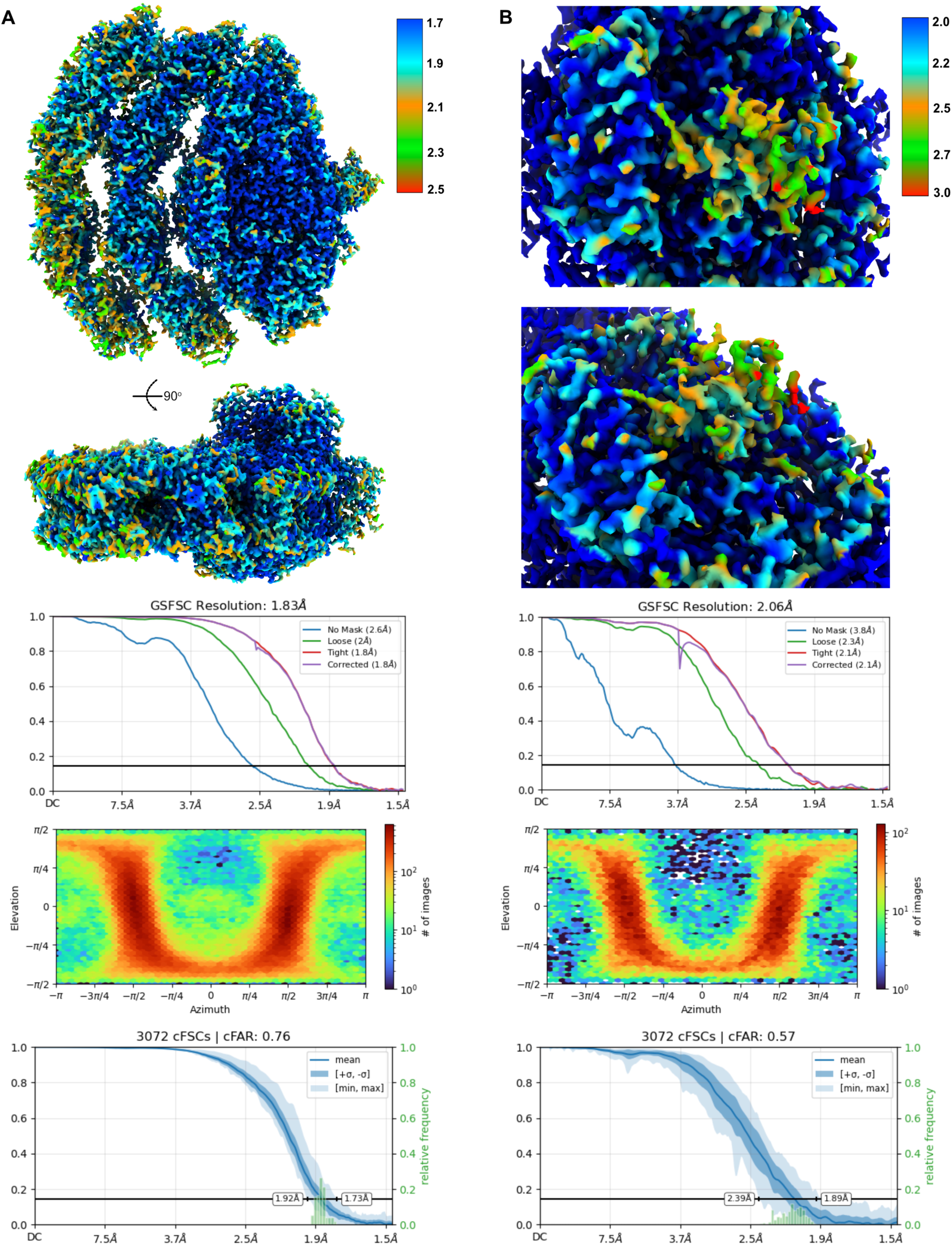
Cryo-EM maps resolution and quality. (A) PSI cryoEM maps colored by their local resolution calculated by GSFSC plot. (B) Cyt *c*_6_-PSI cryoEM maps colored by their local resolution calculated by GSFSC plot.

**Fig. S4.**
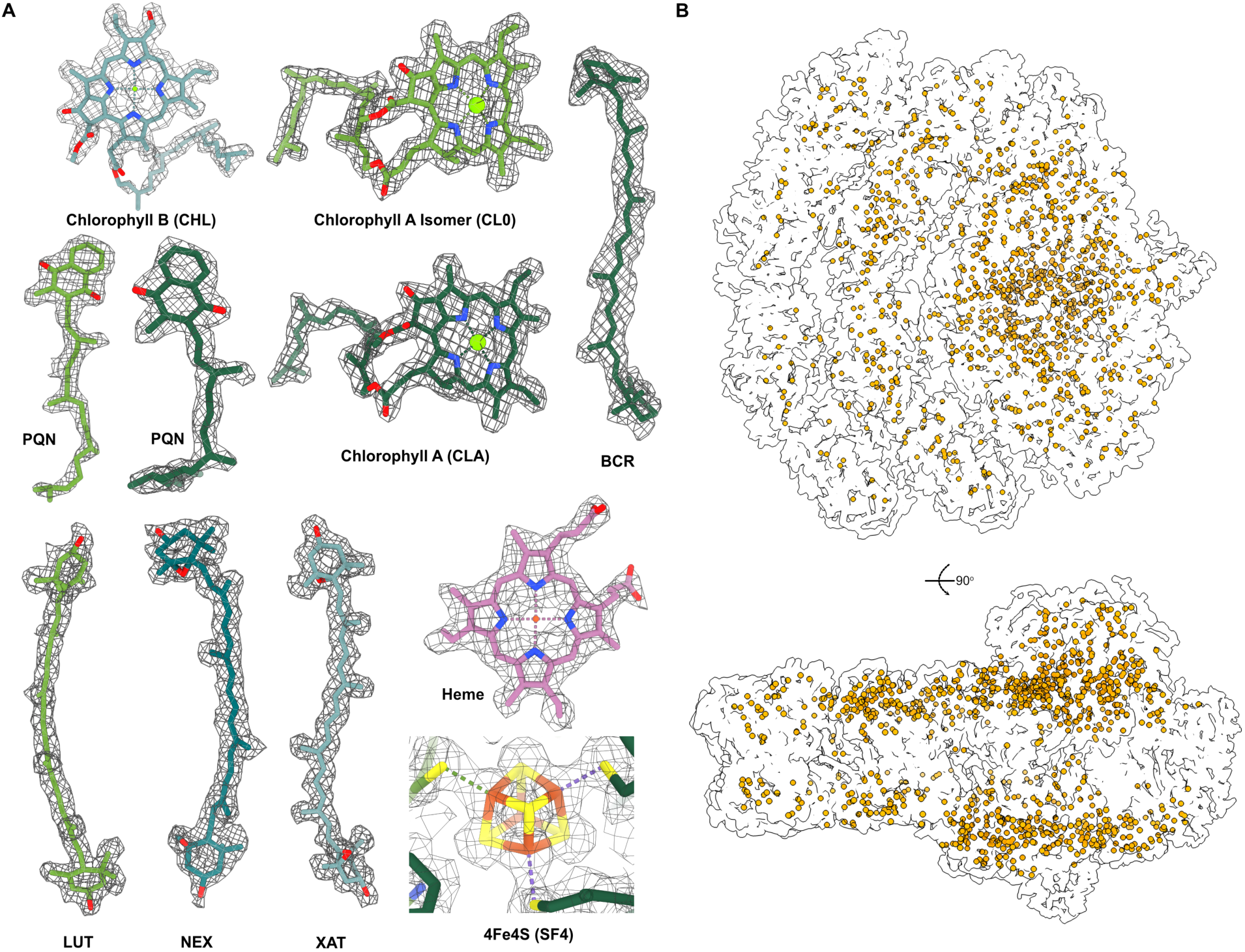
Electron-density maps of co-factors and water molecules distribution. (A) Electron density maps of the co-factors: chlorophyll B (CHL), chlorophyll A isomer (CL0), chlorophyll A (CLA), Phylloquinone (PQN), Beta-Carotene (BCR), (3r,3’r,6s)-4,5-didehydro-5,6-dihydro-beta,beta-carotene-3,3’-diol (LUT), (1r,3r)-6-{(3e,5e,7e,9e,11e,13e,15e,17e)-18-[(1s,4r,6r)-4-hydroxy-2,2,6-trimethyl-7-oxabicyclo[4.1.0]hept-1-yl]-3,7,12,16 tetramethyloctadeca-1,3,5,7,9,11,13,15,17-nonaenylidene}-1,5,5-trimethylcyclohexane-1,3-diol (NEX), (3s,5r,6s,3’s,5’r,6’s)-5,6,5’,6’-diepoxy-5,6,5’,6’-tetrahydro-beta,beta-carotene-3,3’-diol (XAT), 4Fe4S iron/sulfur cluster (SF4). (B) distribution of all the water molecules in orange.

**Fig. S5.**
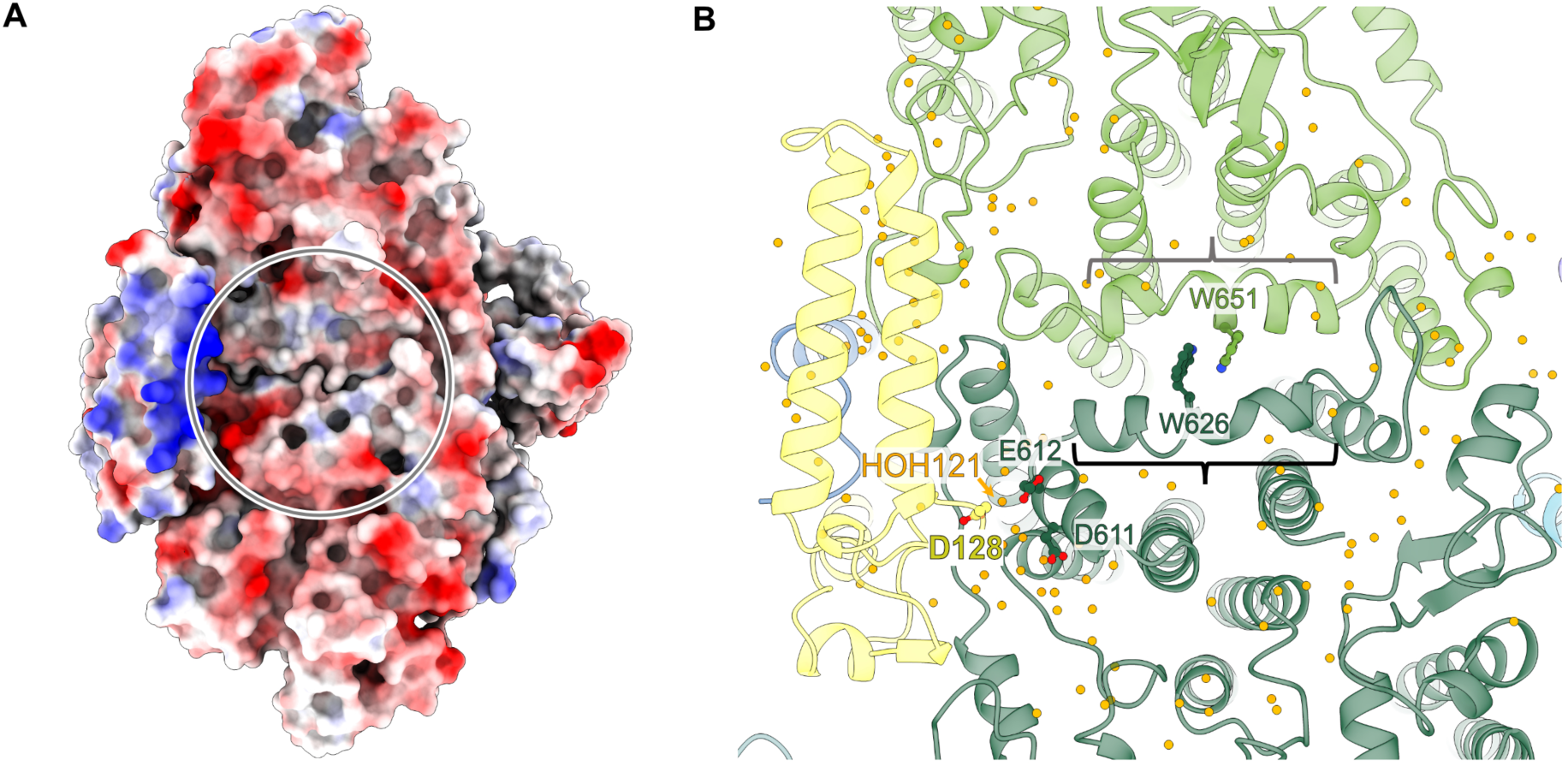
Luminal view of PSI core. (A) Surface electrostatic potential distribution. Positively and negatively charged area are in blue and red, respectively. Gray circle represents the shallow pocket. (B) Cartoon representation with water molecules (orange dots). The Trp dimer is shown. Gray and black brackets indicate PsaA-*l* loop and PsaB-*l’* loop, respectively.

**Fig. S6.**
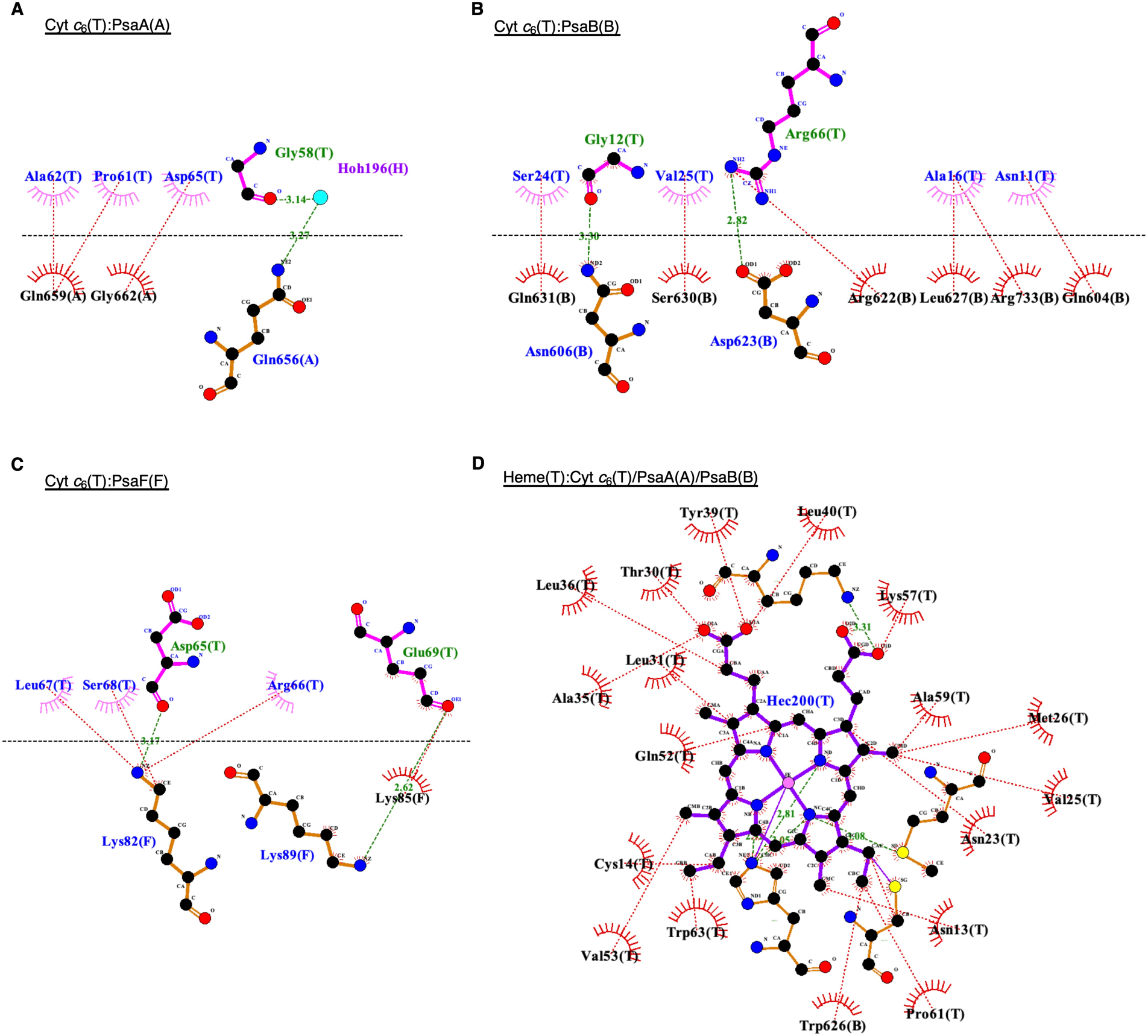
Protein(ligand)-protein interface diagrams of Cyt *c*_6_: PsaA(A), Cyt *c*_6_:PsaB (B) Cyt *c*_6_:PsaF (C) and Heme: Cyt *c*_6_/PsaA/PsaB (D) produced by LigPlot^+^. Green dashed lines with numbers indicate potential hydrogen bonds or electrostatic interactions shorter than 3.35Å, providing the atom-atom distances (Å). Red dashed lines represent other non-bonded contacts including hydrophobic interactions.

**Fig. S7.**
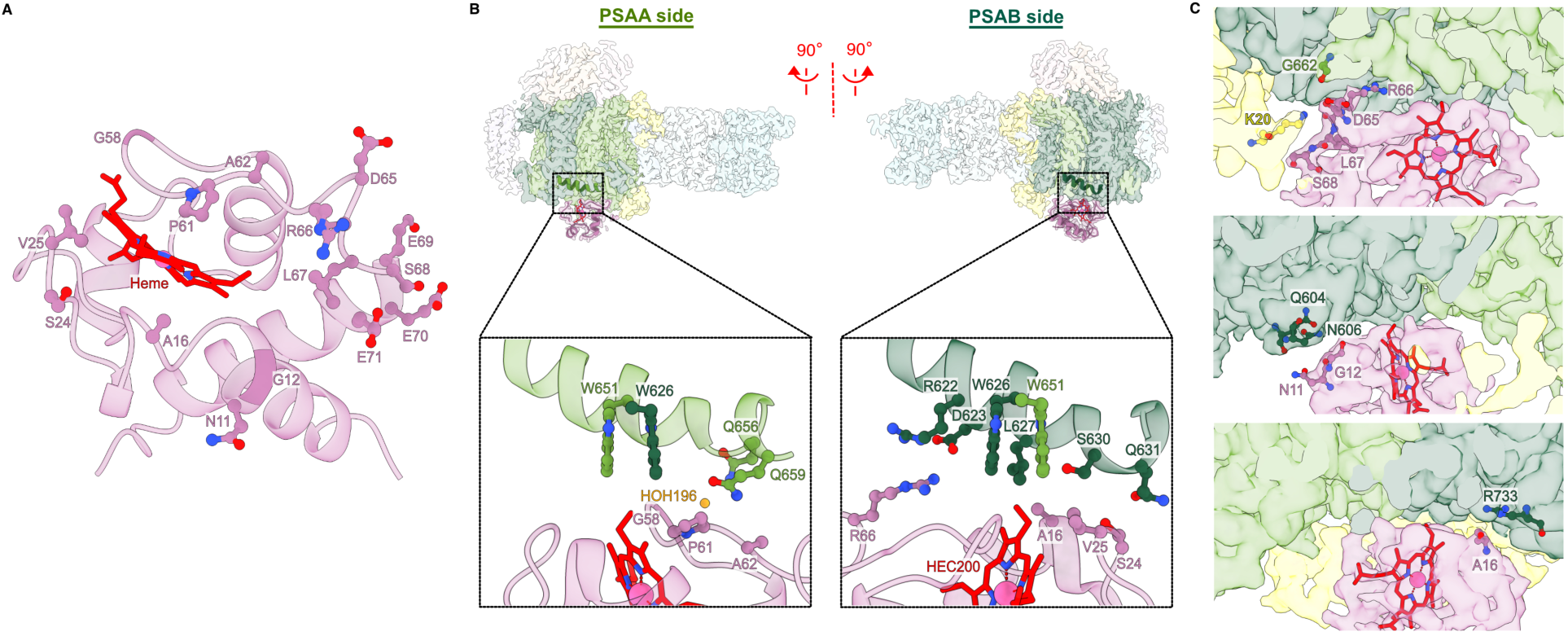
Additional figures for Cyt *c*_6_:PSI binding (A) Stromal view of Cyt *c*_6_:PSI interface (just Cyt *c*_6_ side). Directly interacting amino acids are shown, in addition to E70 and E71. (B) Interfaces between Cyt *c*_6_ and PsaA-*l* loop (left)/PsaB-*l’* loop (right). Directly interacting residues and water molecules (orange) are shown in bottom panels, in addition to PsaA-W651. The Trp dimer is shown in both bottom panels. (C) Peripheral protein-protein interactions on the PsaA-PsaB shallow pocket.

**Fig. S8.**
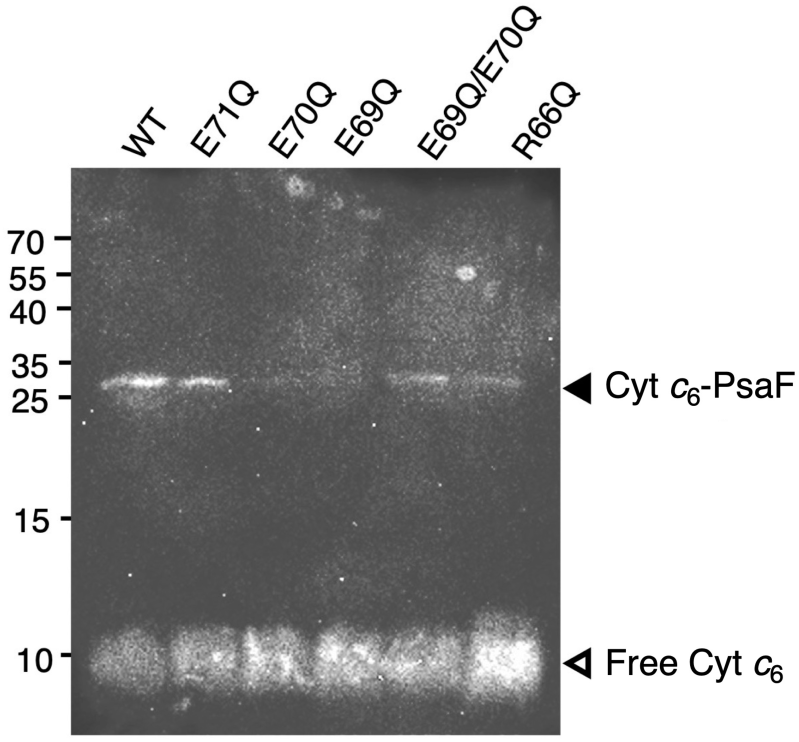
Cross-linking using Cyt *c*_6_ variants with mutations at putatively interacting amino acid residues. The cross-linked samples were subjected to SDS-PAGE and Western blotting.

**Table S1.**
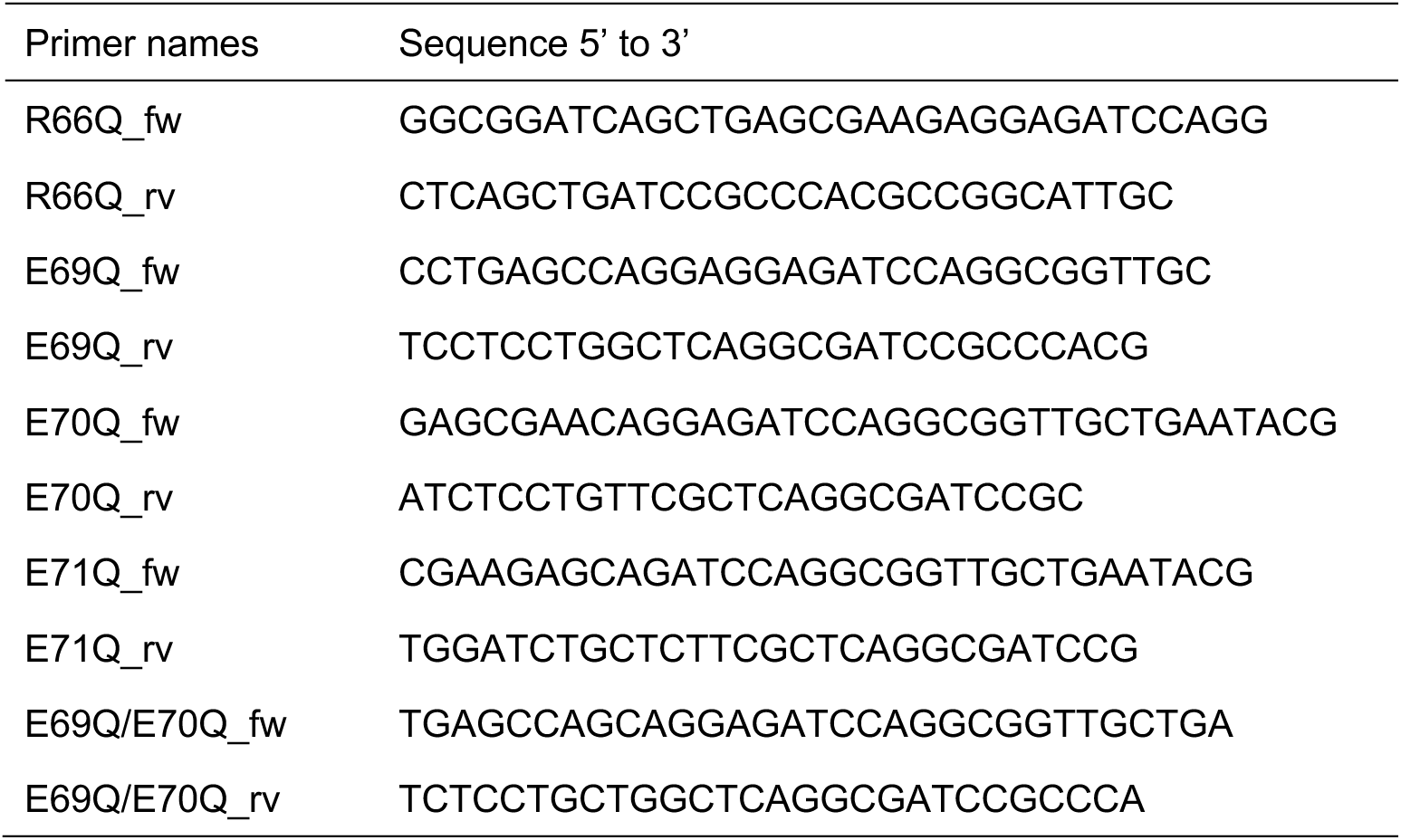
Primers used in this study.

Data S1. (separate file)

Raw data for P700^+^ measured in JTS, including full analysis to the results presented in Fig. 2F

## References

1. C. Castell, L. A. Rodríguez-Lumbreras, M. Hervás, J. Fernández-Recio, J. A. Navarro, New Insights into the Evolution of the Electron Transfer from Cytochrome f to Photosystem I in the Green and Red Branches of Photosynthetic Eukaryotes. Plant Cell Physiol 62, 1082–1093 (2021).

2. M. A. De la Rosa et al., An evolutionary analysis of the reaction mechanisms of photosystem I reduction by cytochrome c(6) and plastocyanin. Bioelectrochemistry 55, 41–45 (2002).

3. G. Sandmann, H. Reck, P. Kessler, P. Boeger. (Arch. Microbiol., 1983), vol. 134, pp. 23–27.

4. A. Díaz-Quintana et al., A comparative structural and functional analysis of cyanobacterial plastocyanin and cytochrome c (6) as alternative electron donors to Photosystem I. Photosynth Res 75, 97–110 (2003).

5. M. Hervas, J. A. Navarro, A. Diaz, H. Bottin, M. A. De la Rosa, Laser-flash kinetic analysis of the fast electron transfer from plastocyanin and cytochrome c6 to photosystem I. Experimental evidence on the evolution of the reaction mechanism. Biochemistry 34, 11321–11326 (1995).

6. F. Drepper, M. Hippler, W. Nitschke, W. Haehnel, Binding dynamics and electron transfer between plastocyanin and photosystem I. Biochemistry 35, 1282–1295 (1996).

7. M. Nordling, K. Sigfridsson, S. Young, L. G. Lundberg, O. Hansson, Flash-photolysis studies of the electron transfer from genetically modified spinach plastocyanin to photosystem I. FEBS Lett 291, 327–330 (1991).

8. W. Haehnel et al., Electron transfer from plastocyanin to photosystem I. EMBO J 13, 1028–1038 (1994).

9. K. Sigfridsson, S. Young, O. Hansson, Structural dynamics in the plastocyanin-photosystem 1 electron-transfer complex as revealed by mutant studies. Biochemistry 35, 1249–1257 (1996).

10. F. Sommer, F. Drepper, M. Hippler, The luminal helix l of PsaB is essential for recognition of plastocyanin or cytochrome c6 and fast electron transfer to photosystem I in Chlamydomonas reinhardtii. J Biol Chem 277, 6573–6581 (2002).

11. M. Hippler, N. Nelson, The Plasticity of Photosystem I. Plant Cell Physiol 62, 1073–1081 (2021).

12. A. Busch, M. Hippler, The structure and function of eukaryotic photosystem I. Biochim Biophys Acta 1807, 864–877 (2011).

13. F. Sommer, F. Drepper, W. Haehnel, M. Hippler, The hydrophobic recognition site formed by residues PsaA-Trp651 and PsaB-Trp627 of photosystem I in Chlamydomonas reinhardtii confers distinct selectivity for binding of plastocyanin and cytochrome c6. J Biol Chem 279, 20009–20017 (2004).

14. I. Díaz-Moreno et al., NMR analysis of the transient complex between membrane photosystem I and soluble cytochrome c6. J Biol Chem 280, 7925–7931 (2005).

15. S. Young, K. Sigfridsson, K. Olesen, O. Hansson, The involvement of the two acidic patches of spinach plastocyanin in the reaction with photosystem I. Biochim Biophys Acta 1322, 106–114 (1997).

16. K. Olesen, M. Ejdebäck, M. M. Crnogorac, N. M. Kostić, O. Hansson, Electron transfer to photosystem 1 from spinach plastocyanin mutated in the small acidic patch: ionic strength dependence of kinetics and comparison of mechanistic models. Biochemistry 38, 16695–16705 (1999).

17. M. Hippler et al., The plastocyanin binding domain of photosystem I. EMBO J 15, 6374–6384 (1996).

18. M. Hippler, F. Drepper, J. Farah, J. D. Rochaix, Fast electron transfer from cytochrome c6 and plastocyanin to photosystem I of Chlamydomonas reinhardtii requires PsaF. Biochemistry 36, 6343–6349 (1997).

19. M. Hippler, F. Drepper, W. Haehnel, J. D. Rochaix, The N-terminal domain of PsaF: precise recognition site for binding and fast electron transfer from cytochrome c6 and plastocyanin to photosystem I of Chlamydomonas reinhardtii. Proc Natl Acad Sci U S A 95, 7339–7344 (1998).

20. M. Hippler, F. Drepper, J. D. Rochaix, U. Mühlenhoff, Insertion of the N-terminal part of PsaF from Chlamydomonas reinhardtii into photosystem I from Synechococcus elongatus enables efficient binding of algal plastocyanin and cytochrome c6. J Biol Chem 274, 4180–4188 (1999).

21. I. Caspy, A. Borovikova-Sheinker, D. Klaiman, Y. Shkolnisky, N. Nelson, The structure of a triple complex of plant photosystem I with ferredoxin and plastocyanin. Nat Plants 6, 1300–1305 (2020).

22. I. Caspy et al., Structure of plant photosystem I-plastocyanin complex reveals strong hydrophobic interactions. Biochem J 478, 2371–2384 (2021).

23. A. Naschberger et al., Algal photosystem I dimer and high-resolution model of PSI-plastocyanin complex. Nat Plants 8, 1191–1201 (2022).

24. J. Li et al., Structure of cyanobacterial photosystem I complexed with ferredoxin at 1.97 Å resolution. Commun Biol 5, 951 (2022).

25. A. Kölsch et al., Current limits of structural biology: The transient interaction between cytochrome. Curr Res Struct Biol 2, 171–179 (2020).

26. F. Sommer, F. Drepper, W. Haehnel, M. Hippler, Identification of Precise Electrostatic Recognition Sites between Cytochrome c6 and the Photosystem I Subunit PsaF Using Mass Spectrometry. J Biol Chem 281, 35097–35103 (2006).

27. B. De la Cerda, A. Díaz-Quintana, J. A. Navarro, M. Hervás, M. A. De la Rosa, Site-directed mutagenesis of cytochrome c6 from Synechocystis sp. PCC 6803. The heme protein possesses a negatively charged area that may be isofunctional with the acidic patch of plastocyanin. J Biol Chem 274, 13292–13297 (1999).

28. F. P. Molina-Heredia, M. Hervás, J. A. Navarro, M. A. De la Rosa, A single arginyl residue in plastocyanin and in cytochrome c(6) from the cyanobacterium Anabaena sp. PCC 7119 is required for efficient reduction of photosystem I. J Biol Chem 276, 601–605 (2001).

29. T. Schwartz, M. Fadeeva, D. Klaiman, N. Nelson, Structure of Photosystem I Supercomplex Isolated from a. Biomolecules 13, (2023).

30. Z. Huang et al., Structure of photosystem I-LHCI-LHCII from the green alga Chlamydomonas reinhardtii in State 2. Nat Commun 12, 1100 (2021).

31. M. Suga et al., Structure of the green algal photosystem I supercomplex with a decameric light-harvesting complex I. Nat Plants 5, 626–636 (2019).

32. I. Caspy et al., Dimeric and high-resolution structures of Chlamydomonas Photosystem I from a temperature-sensitive Photosystem II mutant. Commun Biol 4, 1380 (2021).

33. A. Punjani, J. L. Rubinstein, D. J. Fleet, M. A. Brubaker, cryoSPARC: algorithms for rapid unsupervised cryo-EM structure determination. Nat Methods 14, 290–296 (2017).

34. A. Punjani, H. Zhang, D. J. Fleet, Non-uniform refinement: adaptive regularization improves single-particle cryo-EM reconstruction. Nat Methods 17, 1214–1221 (2020).

35. C. A. Kerfeld, H. P. Anwar, R. Interrante, S. Merchant, T. O. Yeates, The structure of chloroplast cytochrome c6 at 1.9 A resolution: evidence for functional oligomerization. J Mol Biol 250, 627–647 (1995).

36. C. Gerle et al., Three structures of PSI-LHCI from Chlamydomonas reinhardtii suggest a resting state re-activated by ferredoxin. Biochim Biophys Acta Bioenerg 1864, 148986 (2023).

37. Y. Milrad et al., Insights into plastocyanin-cytochrome b6f complex formation: The role of plastocyanin phosphorylation. Plant Physiol 198, (2025).

38. A. C. Wallace, R. A. Laskowski, J. M. Thornton, LIGPLOT: a program to generate schematic diagrams of protein-ligand interactions. Protein Eng 8, 127–134 (1995).

39. R. A. Laskowski, M. B. Swindells, LigPlot+: multiple ligand-protein interaction diagrams for drug discovery. J Chem Inf Model 51, 2778–2786 (2011).

40. S. Kumar, R. Nussinov, Close-range electrostatic interactions in proteins. Chembiochem 3, 604–617 (2002).

41. P. Chakrabarti, J. Janin, Dissecting protein-protein recognition sites. Proteins 47, 334–343 (2002).

42. G. M. Ullmann, M. Hauswald, A. Jensen, N. M. Kostić, E. W. Knapp, Comparison of the physiologically equivalent proteins cytochrome c6 and plastocyanin on the basis of their electrostatic potentials. Tryptophan 63 in cytochrome c6 may be isofunctional with tyrosine 83 in plastocyanin. Biochemistry 36, 16187–16196 (1997).

43. W. Bialek, S. Krzywda, M. Jaskolski, A. Szczepaniak, Atomic-resolution structure of reduced cyanobacterial cytochrome c6 with an unusual sequence insertion. FEBS J 276, 4426–4436 (2009).

44. P. Bernal-Bayard et al., Interaction of photosystem I from Phaeodactylum tricornutum with plastocyanins as compared with its native cytochrome c6: Reunion with a lost donor. Biochim Biophys Acta 1847, 1549–1559 (2015).

45. A. Kölsch et al., Insights into the binding behavior of native and non-native cytochromes to photosystem I from. J Biol Chem 293, 9090–9100 (2018).

46. B. Zhang et al., A High-Resolution Crystallographic Study of Cytochrome c6: Structural Basis for Electron Transfer in Cyanobacterial Photosynthesis. Int J Mol Sci 26, (2025).

47. T. Ueda et al., Structural basis of efficient electron transport between photosynthetic membrane proteins and plastocyanin in spinach revealed using nuclear magnetic resonance. Plant Cell 24, 4173–4186 (2012).

48. T. Inoue et al., Crystal structure determinations of oxidized and reduced plastocyanin from the cyanobacterium Synechococcus sp. PCC 7942. Biochemistry 38, 6063-6069 (1999).

49. G. Finazzi, F. Sommer, M. Hippler, Release of oxidized plastocyanin from photosystem I limits electron transfer between photosystem I and cytochrome b6f complex in vivo. Proc Natl Acad Sci U S A 102, 7031–7036 (2005).

50. M. Antoshvili, I. Caspy, M. Hippler, N. Nelson, Structure and function of photosystem I in Cyanidioschyzon merolae. Photosynth Res 139, 499–508 (2019).

51. R. J. Porra, W. A. Thompson, P. E. Kriedemann. (Biochimica et Biophysica Acta, 1989), vol. 975, pp. 384–394.

52. E. Arslan, H. Schulz, R. Zufferey, P. Kunzler, L. Thony-Meyer, Overproduction of the Bradyrhizobium japonicum c-type cytochrome subunits of the cbb3 oxidase in Escherichia coli. Biochem Biophys Res Commun 251, 744–747 (1998).

53. P. M. Wood, Interchangeable copper and iron proteins in algal photosynthesis. Studies on plastocyanin and cytochrome c-552 in Chlamydomonas. Eur J Biochem 87, 9-19 (1978).

54. S. Q. Zheng et al., MotionCor2: anisotropic correction of beam-induced motion for improved cryo-electron microscopy. Nat Methods 14, 331–332 (2017).

55. A. Rohou, N. Grigorieff, CTFFIND4: Fast and accurate defocus estimation from electron micrographs. J Struct Biol 192, 216–221 (2015).

56. T. Bepler et al., Positive-unlabeled convolutional neural networks for particle picking in cryo-electron micrographs. Res Comput Mol Biol 10812, 245–247 (2018).

57. T. Bepler et al., Positive-unlabeled convolutional neural networks for particle picking in cryo-electron micrographs. Nat Methods 16, 1153–1160 (2019).

58. T. D. Goddard et al., UCSF ChimeraX: Meeting modern challenges in visualization and analysis. Protein Sci 27, 14–25 (2018).

59. P. Emsley, K. Cowtan, Coot: model-building tools for molecular graphics. Acta Crystallogr D Biol Crystallogr 60, 2126–2132 (2004).

60. P. Emsley, B. Lohkamp, W. G. Scott, K. Cowtan, Features and development of Coot. Acta Crystallogr D Biol Crystallogr 66, 486–501 (2010).

61. P. V. Afonine et al., Real-space refinement in PHENIX for cryo-EM and crystallography. Acta Crystallogr D Struct Biol 74, 531–544 (2018).

